# Gene III Ergothioneine ameliorates alcoholic fatty liver disease: A dual strategy of accelerated ethanol elimination and reducing oxidative stress

**DOI:** 10.64898/2026.02.14.705887

**Authors:** Wei Ding, Juan Cao, Cong Guo, Xu Li, Wei Liu, Guohua Xiao

## Abstract

**Background:** Alcoholic fatty liver disease (AFLD) is a progressive hepatic pathology triggered by chronic ethanol consumption, serving as the initial stage of severe liver injury. Currently, there are no FDA-approved pharmacological interventions that specifically target alcohol-induced hepatic steatosis or prevent disease progression, highlighting an urgent need for effective preventive strategies. This study evaluated the preventive efficacy and underlying mechanisms of Gene III Ergothioneine (EGT) in a clinically relevant preclinical model.

**Methods:** C57BL/6 mice were randomized into five groups: a Control group, an alcoholic fatty liver Model group, a Positive control group treated with Silybin (100 mg/kg), and three EGT treatment groups (10, 30, and 50 mg/kg). The NIAAA mouse model was utilized to induce alcoholic fatty liver. Various biochemical, histological, and molecular markers were assessed to evaluate liver damage, alcohol metabolism, lipid profiles, oxidative stress, and inflammation.

**Results:** Gene III EGT treatment significantly ameliorated hepatic steatosis and necrosis, as confirmed by H&E and Oil Red O staining. Notably, EGT accelerated alcohol clearance, reducing serum ethanol levels by up to 54.4% in a dose-dependent manner. Furthermore, EGT restored liver function markers (ALT, AST, GGT) and corrected dyslipidemia by lowering TG, TC, and LDL-C while elevating HDL-C. Mechanistically, EGT suppressed pro-inflammatory cytokines (IL-6, IL-1 β) and mitigated oxidative stress by reducing malondialdehyde (MDA) accumulation and restoring superoxide dismutase (SOD) and glutathione peroxidase (GSH-Px) activities.

**Conclusion:** Gene III Ergothioneine prevents alcoholic liver injury through a dual mechanism: accelerating ethanol metabolism and enhancing hepatocyte antioxidative and anti-inflammatory defenses. These findings position EGT as a promising therapeutic candidate for AFLD management.

## Introduction

Alcoholic fatty liver disease represents the earliest and most prevalent stage of alcoholic liver disease, characterized by excessive triglyceride accumulation in the liver as a direct consequence of chronic ethanol abuse [1]. The pathogenesis of AFLD involves complex mechanisms driven primarily by oxidative stress and disruptions of normal lipid metabolism, which can result in liver damage [2]. Specifically, ethanol metabolism induces the overexpression of cytochrome P450 2E1 and activates NADPH oxidase, leading to excessive production of reactive oxygen species and subsequent depletion of glutathione [3-4]. Furthermore, alcohol-induced gut dysbiosis increases intestinal permeability, allowing endotoxins to reach the liver and activate Kupffer cells, thereby triggering an inflammatory cascade dominated by cytokines such as IL-6, IL-1β, and TNF-α [5]. This imbalance precipitates hepatic oxidative stress, triggering lipid peroxidation, mitochondrial dysfunction, and inflammatory responses that ultimately drive hepatocyte apoptosis and necrosis [6]. While simple steatosis is often considered reversible, it sensitizes the liver to “second hits” such as oxidative stress and inflammation, potentially progressing to steatohepatitis, fibrosis, and cirrhosis if left unchecked.

The global burden of AFLD is substantial, with estimates indicating that alcohol-related harm accounts for over 3 million deaths annually worldwide and contributes to more than 200 health conditions [7]. The current standard of care relies on abstinence [8-9] and nutritional support, with pharmacological agents like Silybin used as supportive therapy. However, traditional antioxidants often lack specific transport mechanisms to achieve high intracellular concentrations in mitochondria, the primary site of ROS Generation.

Ergothioneine, a naturally occurring thiol-histidine derivative, has emerged as a potent dietary antioxidant distinguished by its unique transport system and capacity to mitigate cellular oxidative damage and inflammation [10]. Recent studies have highlighted EGT’s potential in metabolic dysfunction-associated steatotic liver disease (MASLD) via autophagy enhancement and ferroptosis inhibition [11-12].Despite its established antioxidant properties, the specific efficacy of Gene III EGT in preventing alcoholic fatty liver, particularly within the rigorous NIAAA mouse model framework, has not been thoroughly investigated.

Therefore, this study aims to evaluate the preventive effects of Gene III EGT on ethanol-induced hepatic steatosis and injury by comparing its efficacy against Silybin, a standard hepatoprotective agent, in a well-established mouse model of AFLD. Consequently, elucidating the mechanisms by which Gene III EGT modulates alcohol metabolism, lipid profiles, and inflammatory cytokines may provide critical insights for developing novel therapeutic strategies for AFLD.

## 2. Materials and Methods

### 2.1 Chemicals and Reagents

Gene III Ergothioneine (>99.99% purity) was supplied by Jiangsu Gene III Biotechnology Co., Ltd. (Nanjing, China). Silybin (>98.76%) was obtained from Shandong Sikejie Biotechnology Co., Ltd. The Lieber-DeCarli ethanol and control liquid diets were purchased from Xietong Bio (China). Diagnostic kits for ALT, AST, GGT, TG, TC, LDL-C, and HDL-C were obtained from Nanjing Jiancheng Bioengineering Institute (China). ELISA kits for IL-1β, TNF-α, and IL-6 were purchased from Abclonal (China). Kits for GSH-Px, MDA, and Ethanol detection were sourced from Beyotime (China), and the SOD kit was obtained from Addison (China).

### 2.2 Animals and Experimental

Design Specific Pathogen Free (SPF) male C57BL/6 mice (8 weeks old) were obtained from Jiangsu Huachuang Xinnuo Pharmaceutical Technology Co., Ltd. Mice were housed under standard conditions (24°C–28°C, 12h light/dark cycle). All animal procedures were approved by the Animal Ethics Committee (Approval No.: SYXK-2023-0019).

The NIAAA chronic-plus-binge model was established as previously described. After acclimatization, mice were randomized into six groups (n=10/group): (1) Control, (2) Model (ETOH), (3) Positive Control (Silybin 100 mg/kg), and (4–6) Gene III EGT treatment groups (10, 30, and 50 mg/kg). Pretreatment with EGT or Silybin was administered intragastrical for 10 days. Subsequently, ethanol groups were fed the Lieber-DeCarli ethanol liquid diet (5% ethanol) for 10 days, while the Control group received an isocaloric control diet. On day 11, ethanol groups received a single binge of ethanol (5 g/kg), whereas controls received maltodextrin. Mice were sacrificed 9 hours post-binge to collect serum and liver tissues.

### 2.3 Histopathological Analysis

Liver tissues were fixed in 4% paraformaldehyde, embedded in paraffin, and sectioned (5 μm) for Hematoxylin and Eosin (H&E) staining. For lipid droplet visualization, frozen liver sections embedded in OCT compound were subjected to Oil Red O (ORO) staining. Images were captured using a light microscope.

### 2.4 Biochemical Analysis

Serum samples were separated by centrifugation at 4000 rpm for 10 min at 4°C. The levels of liver injury markers (ALT, AST, GGT), lipid profiles (TG, TC, LDL-C, HDL-C), and serum ethanol concentration were quantified using commercial assay kits strictly according to the manufacturers’ instructions. Absorbance was measured using a microplate reader.

### 2.5 Inflammatory and Oxidative Stress Assessment

Serum levels of inflammatory cytokines (IL-6, IL-1β, TNF-α) were determined using enzyme-linked immunosorbent assay (ELISA) kits following the standard protocols. Oxidative stress markers, including MDA content and the activities of SOD and GSH-Px, were assessed using specific colorimetric assay kits according to the manufacturers’ procedures.

### 2.6 Statistical Analysis

Data are expressed as mean ± standard deviation (SD). Statistical differences were analyzed using GraphPad Prism software via one-way analysis of variance (ANOVA) followed by Bonferroni’s post hoc test. A P-value < 0.05 was considered statistically significant

## 3. Results

### 3.1 Effect of Gene III EGT on Liver Histology in NIAAA Mice

To investigate the protective effect of Gene III EGT, we established an NIAAA mouse model. Morphologically, Gene III EGT treatment improved the appearance of fatty livers (Fig. 1A). Histopathological analysis (H&E) revealed that the Control group exhibited normal hepatic cord structure with orderly arranged hepatocytes. In contrast, the Model group showed varying sizes of lipid droplets (black arrows), inflammatory cell infiltration (yellow arrows), and hepatocyte necrosis (red arrows). Compared to the Model group, mice treated with Gene III EGT showed more regular hepatocyte morphology, significantly reduced lipid droplets, and alleviated inflammatory infiltration and pathological injury (Fig. 1B). Consistently, Oil Red O staining demonstrated massive red lipid droplet accumulation in the Model group, which was significantly attenuated by Gene III EGT administration (Fig. 1B). These results confirm that Gene III EGT alleviates alcoholic fatty liver in mice.

**Figure 1.**
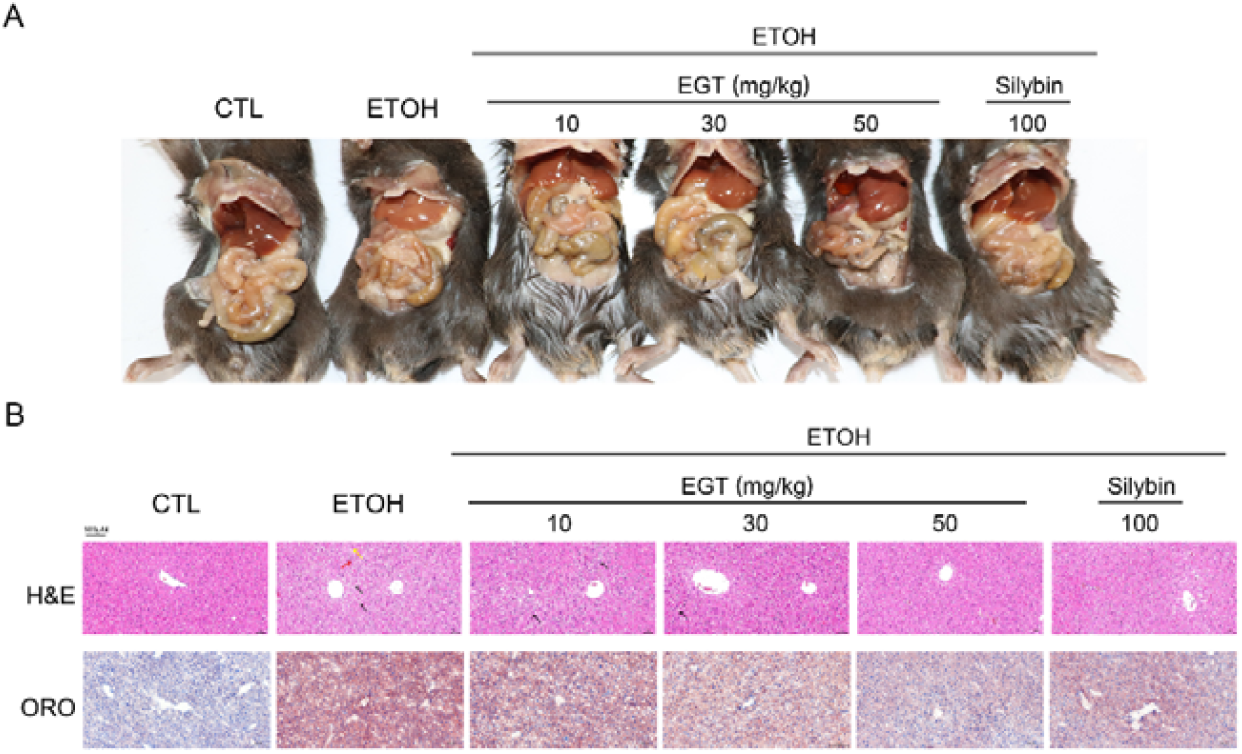
Effect of Gene III EGT on Liver Tissue of NIAAA Mice A mouse alcoholic fatty liver model was established to verify the effect of Gene III EGT on mouse liver tissue. A. Mouse anatomy diagram. B. Representative photos of H&E sections and Oil Red O staining of mouse liver tissue (200x).

### 3.2 Effect of Gene III EGT on Serum Ethanol Levels

As shown in Fig. 2, serum ethanol levels were significantly elevated in the Model group compared to the Control group. Gene III EGT administration reduced serum ethanol levels by 20.4%, 45.2%, and 54.4% in the low, medium, and high-dose groups, respectively, compared to the Model group, exhibiting a dose-dependent trend. This suggests that Gene III EGT promotes alcohol metabolism, thereby lowering serum alcohol content. And this magnitude of reduction significantly exceeded that of Silybin (∼30% reduction), suggesting that EGT may actively enhance the enzymatic clearance of ethanol or protect metabolic enzymes from inactivation.

**Figure 2.**
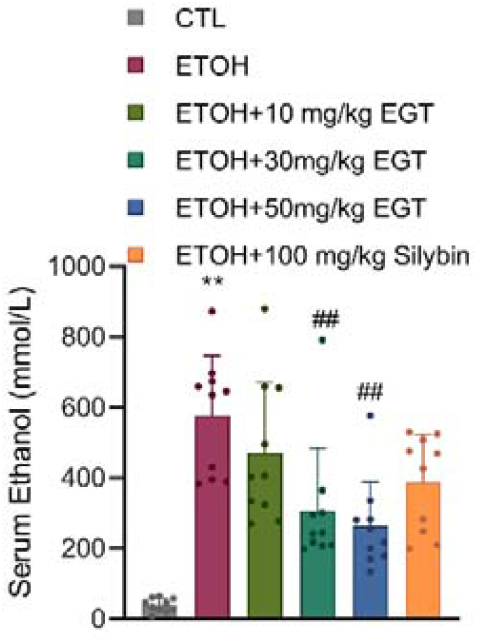
Effect of Gene III EGT on Serum Alcohol Content in NIAAA Mice Alcohol content in mouse serum. n=10, ** P < 0.01 v.s. CTL group, ## P < 0.01 v.s. ETOH group, data are expressed as Mean±SD.

### 3.3 Effect of Gene III EGT on Liver Injury Markers

Compared to the Control group, the Model group showed significantly increased serum levels of ALT, AST, and GGT. Gene III EGT treatment significantly reduced these markers. Specifically, AST levels decreased by 25.8%, 21.5%, and 50.2% (Fig. 3A); ALT levels decreased by 52.6%, 63.7%, and 72.8% (Fig. 3B); and GGT levels decreased by 9%, 10.4%, and 33% (Fig. 3C) compared to the Model group. These findings confirm the hepatoprotective effect of Gene III EGT.

**Figure 3.**
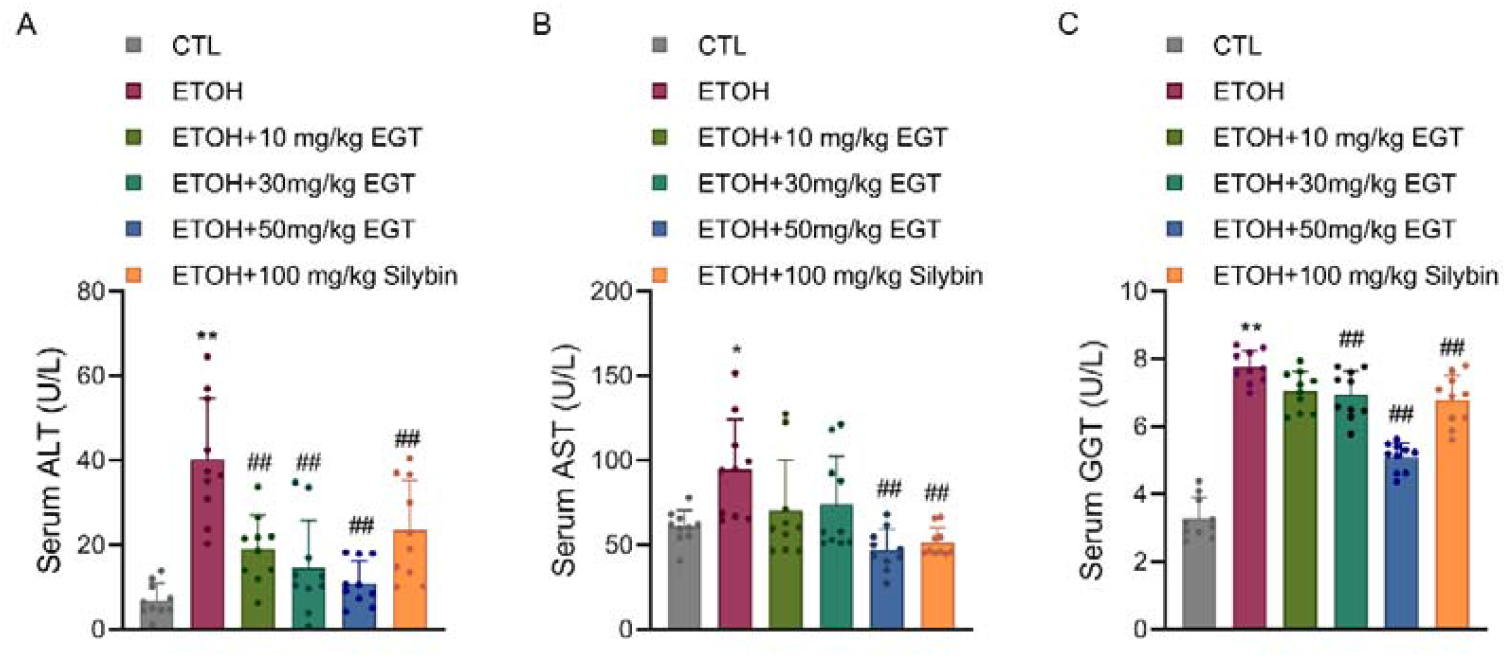
Effect of Gene III EGT on Liver Injury Indicators in NIAAA Mice A. Mouse serum ALT content. B. Mouse serum AST content. C. Mouse serum GGT content. n=10, * P < 0.05, ** P < 0.01 v.s. CTL group, ## P < 0.01 v.s. ETOH group, data are expressed as Mean±SD.

### 3.4 Effect of Gene III EGT on Serum Lipid Profiles

Regarding lipid biochemical indicators, serum TG and TC levels were significantly elevated in the Model group compared to the Control group. Gene III EGT treatment reduced TG levels by 10.6%, 20.3%, and 40.6%, and TC levels by 17.6%, 31.4%, and 40.4%, respectively (Fig. 4A-B). Additionally, LDL-C levels, which were elevated in the Model group, decreased by 9.3%, 8.1%, and 22.9% following EGT treatment (Fig. 4C). Conversely, HDL-C levels, which showed a decreasing trend in the Model group, increased by 20.96%, 35%, and 31.4% with EGT treatment (Fig. 4D). This indicates that Gene III EGT promotes the restoration of normal serum lipid profiles.

**Figure 4.**
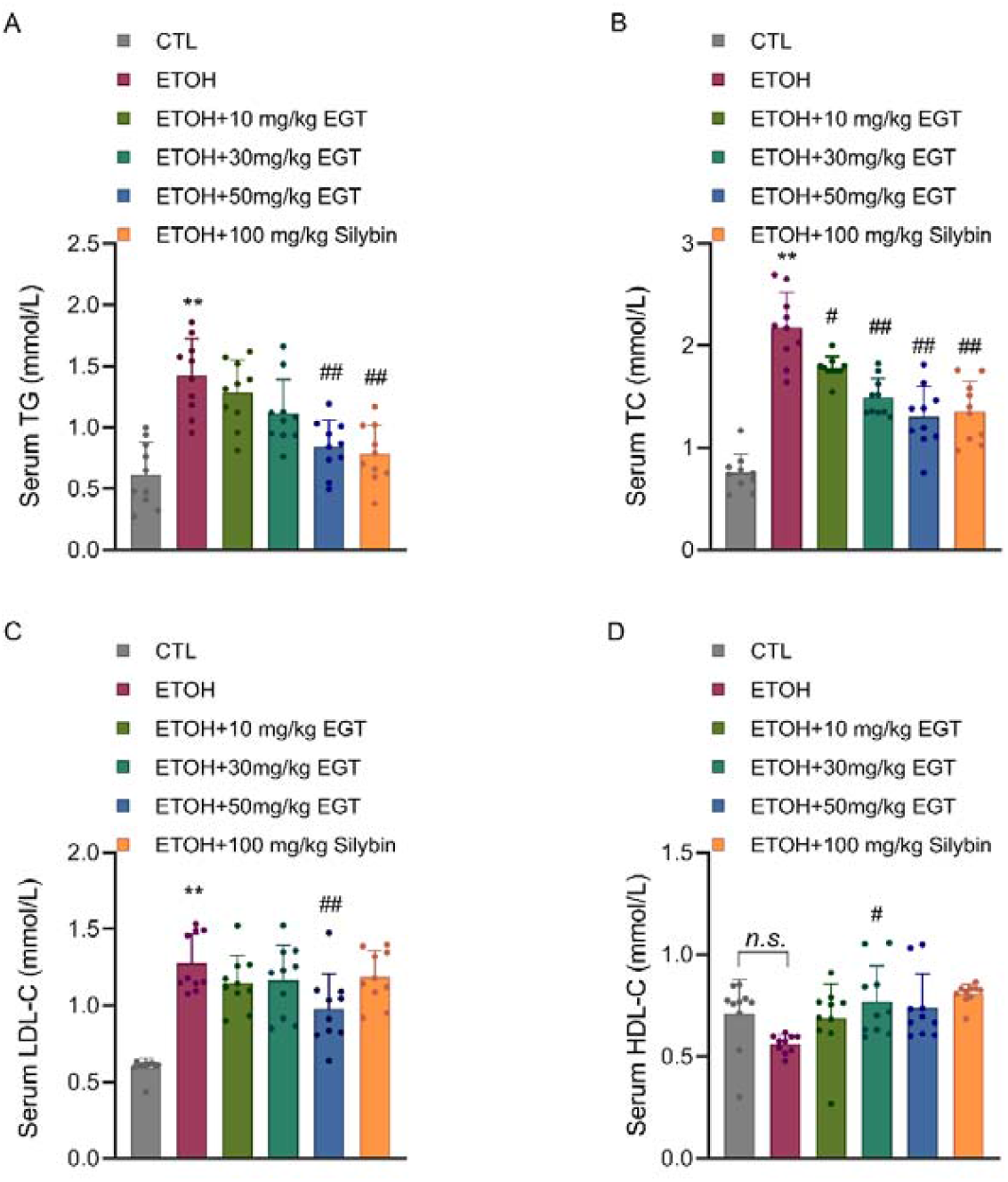
Effect of Gene III EGT on Blood Lipid Content in NIAAA Mice A. Mouse serum TG content. B. Mouse serum TC content. C. Mouse serum LDL-C content. D. Mouse serum HDL-C content. n=10, ** P < 0.01 v.s. CTL group, # P < 0.05, ## P < 0.01 v.s. ETOH group, n.s.: no significance, data are expressed as Mean±SD.

### 3.5 Effect of Gene III EGT on Serum Inflammatory

Cytokines In terms of inflammation, the Model group exhibited significantly elevated levels of pro-inflammatory cytokines IL-6 and IL-1β compared to the Control group. Gene III EGT treatment significantly lowered serum IL-6 levels by 17.5%, 33%, and 60.5%, and IL-1β levels by 42.1%, 55.6%, and 67.1%, in a concentration-dependent manner (Fig. 5A-B). Interestingly, TNF-α levels did not show significant variance at the 9-hour post-binge time point, likely due to its transient peak occurring earlier in the inflammatory cascade (Fig. 5C). This suggests that Gene III EGT alleviates the inflammatory state in mice.

**Figure 5.**
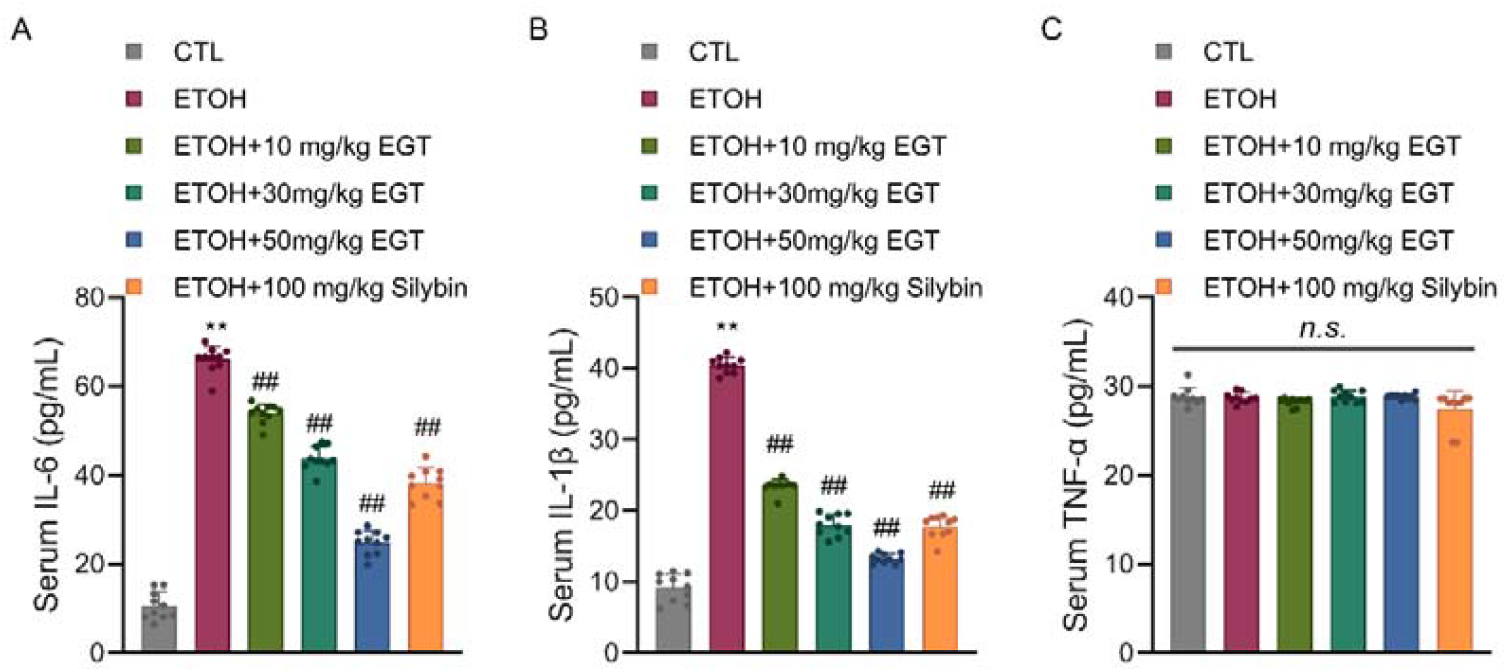
Effect of Gene III EGT on Serum Inflammatory Factor Content in NIAAA Mice Mouse serum IL-6 content. B. Mouse serum IL-1β content. C. Mouse serum TNF-α content. n=10, ** P < 0.01 v.s. CTL group, ## P < 0.01 v.s. ETOH group, n.s.: no significance, data are expressed as Mean±SD.

### 3.6 Effect of Gene III EGT on Oxidative Stress Indices

Regarding oxidative stress, serum MDA levels were significantly increased in the Model group. Gene III EGT treatment reduced MDA levels by 21.5%, 41%, and 50.2% in a concentration-dependent manner (Fig. 6A). Furthermore, the activities of antioxidant enzymes SOD and GSH-Px were significantly decreased in the Model group. Compared to the Model group, EGT treatment increased SOD levels by 16.4%, 62.3%, and 46.7% (Fig. 6B), and GSH-Px activity by 71.4%, 140.2%, and 198.2% (Fig. 6C). These results suggest that GENEIII EGT improves the oxidative stress status in mice.

**Figure 6.**
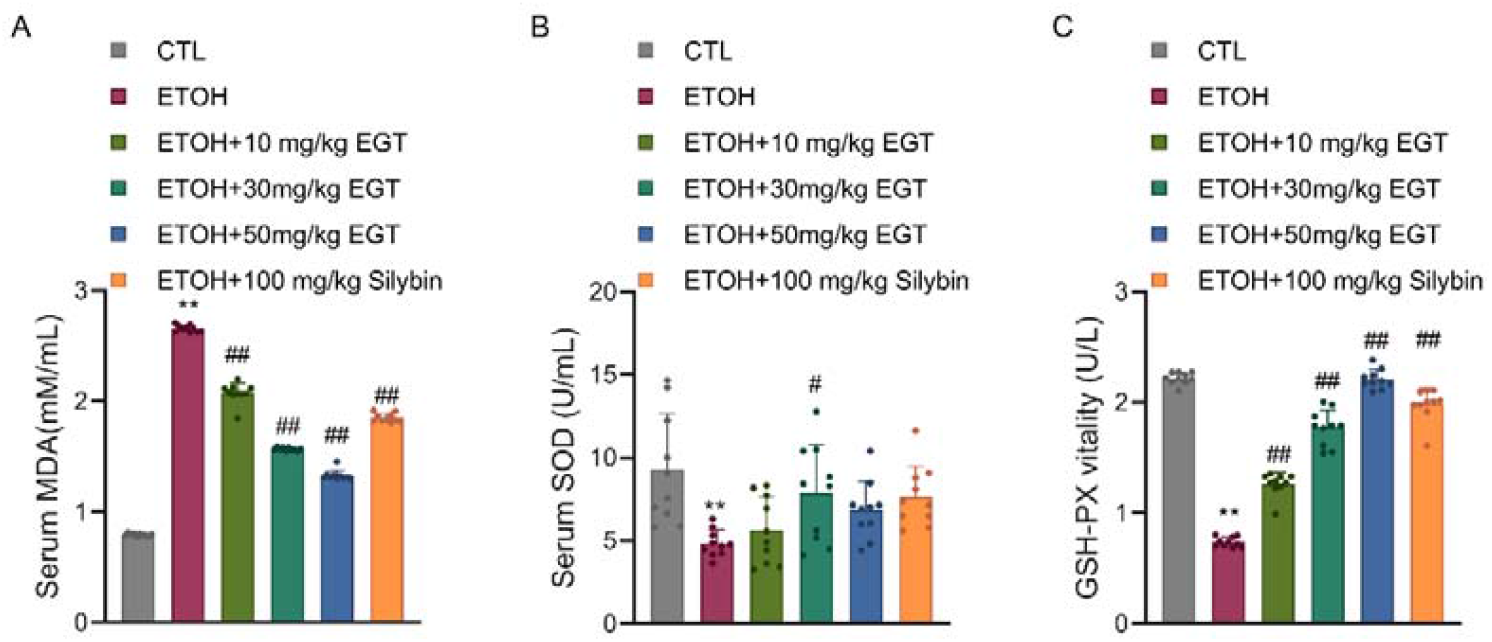
Effect of Gene III EGT on Oxidative Stress Indicators in NIAAA Mice A. Mouse serum MDA content. B. Mouse serum SOD content. C. Mouse serum GSH-PX content. n=10, ** P < 0.01 v.s. CTL group, # P < 0.05, ## P < 0.01 v.s. ETOH group, data are expressed as Mean±SD.

Interestingly, while the high-dose EGT (50 mg/kg) significantly elevated SOD activity compared to the model group (+46.7%), the magnitude of this increase was slightly lower than that of the medium-dose group (+62.3%). This observation may be attributed to a homeostatic feedback mechanism. Given that EGT itself is a potent direct scavenger of reactive oxygen species (ROS), the high intracellular concentration of EGT might have sufficiently reduced the basal oxidative stress signal required to induce maximal SOD expression. Furthermore, considering the continuous and robust surge in GSH-Px activity at the high dose (+198.2%), this pattern suggests a synergistic shift in the antioxidant defense strategy: prioritizing the clearance of downstream peroxides (via GSH-Px) to prevent the accumulation of hydrogen peroxide produced by SOD, thereby optimizing the overall detoxification efficiency.

## Discussion

This study systematically evaluated the preventive efficacy of Gene III EGT on alcoholic fatty liver disease (AFLD) using the NIAAA chronic-plus-binge mouse model. Our findings demonstrate that GENE III EGT not only significantly ameliorates hepatic pathology, reduces liver injury markers, and restores lipid homeostasis but also reveals a unique “dual-defense mechanism”: preventing liver injury through the synergistic action of accelerating ethanol metabolic clearance and enhancing endogenous hepatocellular antioxidant/anti-inflammatory defenses.

The most striking finding of this study is the capacity of Gene III EGT to accelerate alcohol clearance. High-dose Gene III EGT reduced serum ethanol levels by 54.4%, significantly outperforming the standard agent Silybin. We hypothesize that this effect may be attributed to the protection of alcohol dehydrogenase (ADH) and aldehyde dehydrogenase (ALDH) from oxidative inactivation [13]. As a stable thione-thiol antioxidant, EGT may maintain the structural integrity of critical thiol groups within these metabolic enzymes, ensuring efficient alcohol metabolism and thereby reducing the sustained toxicity of ethanol at its source [14].

Regarding oxidative stress, EGT treatment significantly reduced lipid peroxidation (MDA) and restored antioxidant enzyme activities in a distinct synergistic pattern. While SOD activity in the high-dose EGT group was significantly elevated compared to the Model group, the magnitude of induction was slightly lower than in the medium-dose group, showing a non-linear trend. In contrast, GSH-Px activity exhibited a continuous and robust dose-dependent surge (increasing nearly 2-fold). This phenomenon likely reflects a homeostatic feedback regulation of the antioxidant system: the potent direct radical-scavenging ability of high-concentration EGT may reduce the oxidative stress threshold required to induce SOD [15]. Simultaneously, the liver prioritizes a substantial upregulation of the downstream detoxifying enzyme (GSH-Px) to ensure the rapid clearance of hydrogen peroxide Generated by SOD. This “demand-based” enzyme regulation highlights the intelligent characteristic of EGT in maintaining cellular redox balance.

Furthermore, Gene III EGT effectively blocked the inflammatory cascade by inhibiting the release of pro-inflammatory cytokines IL-6 and IL-1β, potentially via the suppression of the NF-κB signaling pathway [16]. In terms of lipid metabolism, EGT not only improved the lipid profile but also significantly elevated HDL-C levels. Given that EGT is specifically accumulated in mitochondria via the OCTN1 transporter [10, 17], it likely maintains fatty acid oxidation efficiency by protecting mitochondrial function and may also facilitate lipophagy to accelerate lipid droplet degradation [11].

In conclusion, Gene III EGT effectively prevents alcoholic liver injury through a dual mechanism of “metabolic enhancement” and “cytoprotection.” These findings provide a solid scientific basis for developing Gene III EGT as a novel functional food or therapeutic agent for the prevention of AFLD.

